# Adolescent sleep is critical for the developmental shaping of social novelty preference

**DOI:** 10.1101/2020.06.29.178350

**Authors:** Wen-Jie Bian, Luis de Lecea

## Abstract

Sleep takes one-third of our lives, yet its functions remain largely unknown. A large proportion of young patients with neurodevelopmental disorders such as autism spectrum disorders (ASDs) and schizophrenia have sleep problems, including delayed sleep onset, shortened sleep duration and sleep fragmentation, which have been linked to social interaction deficit, a shared symptom of these disorders. However, the causal relationship between sleep disruption and social defects as well as the underlying mechanisms have not yet been established despite its importance in understanding the etiology of these disorders and developing potential therapeutic means. Here using the three-chamber social interaction test, we found that developmental sleep disruption (SD) in adolescent mice caused significant and long-lasting impairment in the preference towards social novelty during adult social interactions without affecting the overall sociality. Interestingly, SD performed in the adulthood did not induce any social defect, indicating a critical period within adolescence during which sleep shapes social novelty preference. Furthermore, by analyzing the adolescent sleep and adult social behavior in a mouse model of Shank3 mutation that mimics a genetic aberrance in ASDs, we found that the development of sociality is correlated with adolescent NREM sleep while social novelty preference is correlated with adolescent REM sleep. Collectively, these results demonstrate a critical role of adolescent sleep in the forming of social novelty preference and the developmental shaping of social behavior.

## Introduction

Virtually all higher organisms sleep and in humans, sleep takes approximately one third of our lives(Siegel, 2008). During sleep, the sensory inputs are relatively inhibited, muscle activity is largely reduced, our body is dramatically less reactive to surrounding stimuli and our mind stays unconscious(Krueger et al., 2015). Sleep is believed to be fundamental and essential for the well-being of an organism, serving metabolic needs of the body as well as cognitive and mental purposes of the brain(Frank, 2012; Krueger et al., 2015; Siegel, 2008). Extensive studies have demonstrated a role of sleep in brain functions such as memory consolidation and neural plasticity (Frank, 2012; Krueger et al., 2015; Tononi and Cirelli, 2014), however the underlying mechanisms are largely unknown.

Studies in both human subjects and lab animals have shown that sleep architecture has a clear developmental trajectory. Total sleep time decreases from infancy through adolescence to adulthood (Huber and Born, 2014). In early childhood, electroencephalogram (EEG) hallmarks emerge and gradually correlate with vigilance states, i.e. wakefulness, rapid-eye-movement (REM) sleep and non-REM (NREM) sleep (Cirelli and Tononi, 2015). Additionally, compared with adult sleep, adolescent sleep shows more slow waves and sleep spindles(Campbell and Feinberg, 2009; de Vivo et al., 2014), which are EEG events believed to play important roles in cognitive functions of sleep such as memory consolidation. Furthermore, a large portion (50 – 80%) of young patients of developmental psychiatric disorders including autism spectrum disorders (ASDs) and schizophrenia (SZ) have been reported to have sleep problems, including delayed sleep onset, shortened sleep duration and fragmentation of sleep continuity(Carmassi et al., 2019; Heussler, 2016; Kaskie et al., 2017; Mattai et al., 2006; Robinson-Shelton and Malow, 2016; Veatch et al., 2017). More detailed diagnostic assessments have revealed a significant inverse correlation between sleep duration and severity of social communication deficit, a shared core symptom of these psychiatric disorders(Veatch et al., 2017). In contrast, although association between higher rates of repetitive behaviors (RRB) and increased sleep disturbance has been documented in ASD patients, the correlation between sleep and a specific type of RRB was rather weak (Hundley et al., 2016; Veatch et al., 2017). Collectively, these pieces of evidence indicate that sleep at certain developmental stage may be of particular importance for social behavior.

Neural circuits are more plastic and dynamic throughout postnatal development, and hence more vulnerable, than that in adulthood. Synaptic connections rapidly form in the early postnatal life and are subsequently removed through substantial synapse/spine pruning during adolescence, before their formation and elimination are balanced and the total synapse number becomes stable in the adulthood (Bhatt et al., 2009; Bian et al., 2015; Moyer and Zuo, 2018). Interestingly, several studies have shown that disrupting sleep during certain developmental stages leads to dramatic consequences in the connectivity and functions of brain networks in multiple species. For instance, in the visual cortex of developing cats, sleep deprivation, and especially REM deprivation, abolished the ocular dominance (OD) shift induced by the prior monocular visual experience (Dumoulin Bridi et al., 2015; Frank et al., 2001), while in developing rats, suppression of REM prolonged the time window for cortical long-term potentiation induction (Shaffery et al., 2006; Shaffery et al., 2002). In vivo live imaging using two-photon microscopy and ultrastructural studies have shown that sleep affects synaptic dynamics in both neocortex (de Vivo et al., 2017; Li et al., 2017a; Maret et al., 2011; Yang et al., 2014) and hippocampus(Spano et al., 2019) of juvenile mice, despite whether it promotes synaptic formation/strengthening or elimination/weakening remain controversial. Furthermore, sleep disruption during development impacts courtship behavior in flies, while sleep deprivation at older age does not (Kayser et al., 2014). These results shed light on a sophisticated role of sleep/wake cycle during postnatal development, and particularly during adolescent development. Both NREM and REM sleep impact heavily on neural circuit dynamics and synaptic plasticity during adolescence, but the functional significance at behavioral level and the underlying mechanism remain unclear.

Here we address this question by performing sleep disruption (SD) during critical time window within adolescence and probing the effect of SD on adult social interaction. We further establish associations between abnormal social preference in adulthood and certain sleep components during adolescence.

## Results

### 1. Sleep disruption during adolescence

To study during which developmental stage sleep may play a role for adult social behavior, we first sought to examine how social interaction behavior is shaped during the postnatal development. We started by probing the same-sex social interactions in wildtype (WT), male C57BL6 mice of different ages using the 3-chamber social interaction test. We used the experimental procedure as described elsewhere (Kaidanovich-Beilin et al., 2011) (see Method). In brief, we placed the test mice in an apparatus with three compartment chambers. After habituation, we allowed the test mice a 10-minute trial (Trial 1) to freely explore the three chambers and interact with a never-before-met conspecific (Stranger 1, S1) contained in a mesh cup in one chamber and an empty mesh cup (Empty, E) in the opposite chamber, serving as the non-social novel object. After Trial 1, a second 10-minute trial (Trial 2) began, and a second never-before-met conspecific (Stranger 2, S2) was placed in the previously empty cup, serving as the social novelty stimulus while the Stranger 1 had become familiar. At age of postnatal day 28 (P28), test mice spent almost equal time interacting with E, S1 and S2, and we did not observe any obvious preference in either sociality (Trial1, S1 vs. E) or social novelty (Trial 2, S2 vs. S1, Fig. S1). Both preferences showed trends of gradual increase as the development proceeded from P28 to P56 and became stabilized or even slightly decreased as the animal entered young adulthood (> P56, Fig. S1).

The time window when both sociality preference and social novelty preference demonstrated most dramatic increases (approximately P28 – P56) overlaps mostly with a middle phase of adolescence in mice that is tightly associated with puberty and also termed as “periadolescence” or “pubescent” (P34 – P46)(Brust et al., 2015; Laviola et al., 2003). The neural circuits and synaptic connections in the brain undergo tremendous remodeling and refinement during this period which is thought to be important for adult brain functions and can be dramatically influenced by experiences or environmental stimuli (Bhatt et al., 2009; Bian et al., 2015; Moyer and Zuo, 2018). We surmise that sleep during this developmental time window should play a role in shaping both social preferences. Thus, we sought to perform sleep disruption (SD) in the early light phase of adolescence for 4 hours per day (Zeitgeber time, ZT 2 – 6) and for 5 consecutive days between P35 – P42. The SD was achieved through a programmed apparatus which automatically generated randomized number of pushes (2 – 8) to the bottom of the chamber at randomized intervals (every 0.2 – 0.8 minutes) to keep the animals awake (see Methods). Simultaneous electroencephalogram (EEG) and electromyography (EMG) recordings validated that this protocol abolishes both non-rapid-eye-movement (NREM) and rapid-eye-movement (REM) sleep and causes rebound of both sleep states after daily SD session (Fig. 1A, 1B and 1E–1H). Despite this sleep recovery, there were slight but significant reductions in both NREM and REM sleep and increase in Wake amount across the total 24-hour cycle during the days with SD (Fig. 1I). This automatic SD protocol did not induce any freezing, fighting or other observable behavioral abnormalities when the SD session was over, and the mice were returned to their homecages. We did not detect substantial stress at the end of SD sessions using ELISA of plasma corticosterone (Fig. 1J), despite that chronic subthreshold stress or transient stress during early hours of SD sessions cannot be excluded. This manipulation does not alter sleep architecture permanently, as 24 hours after the final SD day, the sleep pattern has already recovered and indistinguishable from the baseline, and no obvious change was found in the relative power of all frequency bands (Fig. 1A–1I). Control littermates received the same total amount of pushes in the same apparatus expect that it was performed in early dark phase (ZT 12 – 16) and in fixed frequency (control protocol, Ctrl), which presumably had neglectable impact on adolescent sleep.

**Figure 1.**
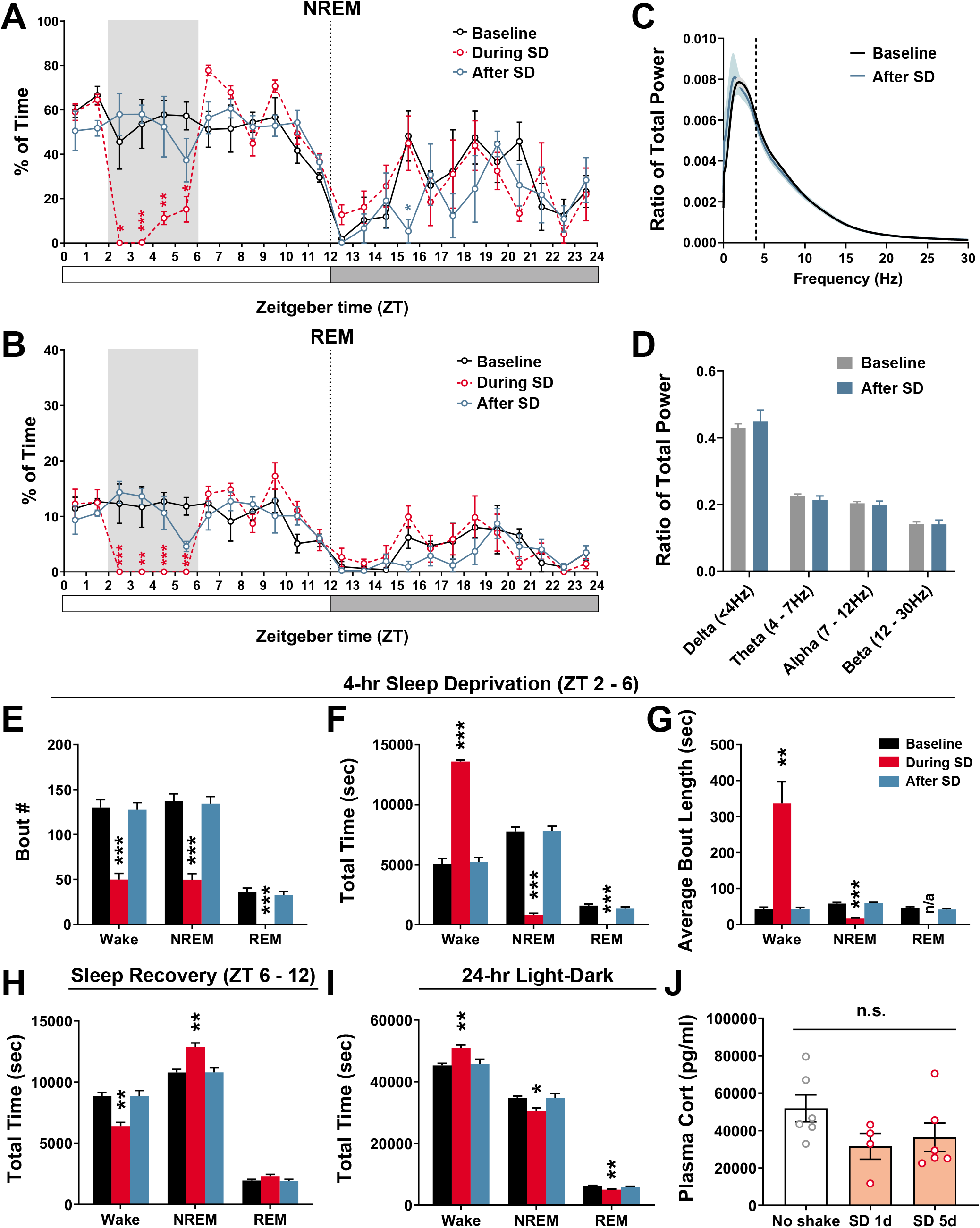
Adolescent sleep disruption. (A, B) The percentage of time that adolescent mice (P35 – 42) spent in NREM (A) and REM (B) sleep in every hour over a 24-hr light-dark cycle before (Baseline, black), during (red), and after (blue) the 5 days containing daily SD sessions (ZT 2 – 6, indicated by the grey stripe). The 3 groups of data were from the same mice (n = 4), and During SD and After SD were compared to Baseline. (C, D) EEG power spectrum of NREM sleep and relative power of each frequency band was not significantly altered after 5 days of SD compared to Baseline (n = 9). (E – G) The bout number (E), total amount (F) and average bout length (G) of Wake, NREM and REM states between ZT 2 – 6 (n = 9). (H, I) The total amount of Wake, NREM and REM during ZT 6 – 12 and over the 24-hr light-dark cycle (n = 9). (J) Plasma corticosterone level of adolescent mice (P35 – 42) after the 1st day and 5 days of SD compared to naïve mice of same age receiving no shake (No shake, n = 6; SD 1d, n = 4; SD 5d, n = 6). All data are shown as mean ± s.e.m. The *p*-values were calculated using statistical tests listed in Supplementary Table 1. * *p* < 0.05; ** *p* < 0.01; *** *p* < 0.001; n.s., not significant.

### 2. Adolescent sleep disruption impairs social novelty preference

After receiving 5 days of SD or Ctrl protocols between P35 – P42, mice were left undisturbed for 2 weeks and tested for social preferences using the 3-chamber assay at P56 – P60. As shown in Fig. 2A, Ctrl mice developed strong sociality preference, as suggested by more interactions with the social object (S1) compared to the non-social object (the empty cup, E). Ctrl mice also showed a strong preference towards social novelty, as shown by the time they spent interacting with the novel stimulus mouse (S2) compared with the familiar one (S1) in Trial 2 (Fig. 2A). However, mice that went through SD in their adolescence showed no clear preference during the social novelty trial spending almost equal time with both stimulus mice, suggesting impaired social novelty preference, although their sociality preference seemed to be intact (Fig. 2A). In all experiments, the preference index of social novelty in SD group was significantly lower than that in Ctrl group and close to zero, while the preference index of sociality showed no difference between the groups (Fig. 2B). These effects of adolescent SD were similar between using the automatic system and with the manual, “gentle-touch” protocol (Fig. S2A, see Methods). Thus, all following experiments were performed using the automatic SD system. We also performed SD at a later adolescent stage of P42 – 49, which led to similar defect in social novelty preference but not in sociality preference (Fig. S2B). Additionally, both Ctrl and SD mice showed similar numbers of crossing doors between the chambers during the whole test session (Fig. 2C), suggesting that overall locomotor activity was not changed by adolescent SD. Elevated plus maze test also suggests that adolescent SD did not increase anxiety during young adulthood when the social test was performed (Fig. S2C). In addition to social novelty impairment, the SD mice also showed reduced interaction time with novel, nonsocial objects in the novel object recognition test with reduced memory component (see Methods), suggesting their preference towards non-social novelty is also impaired.

**Figure 2.**
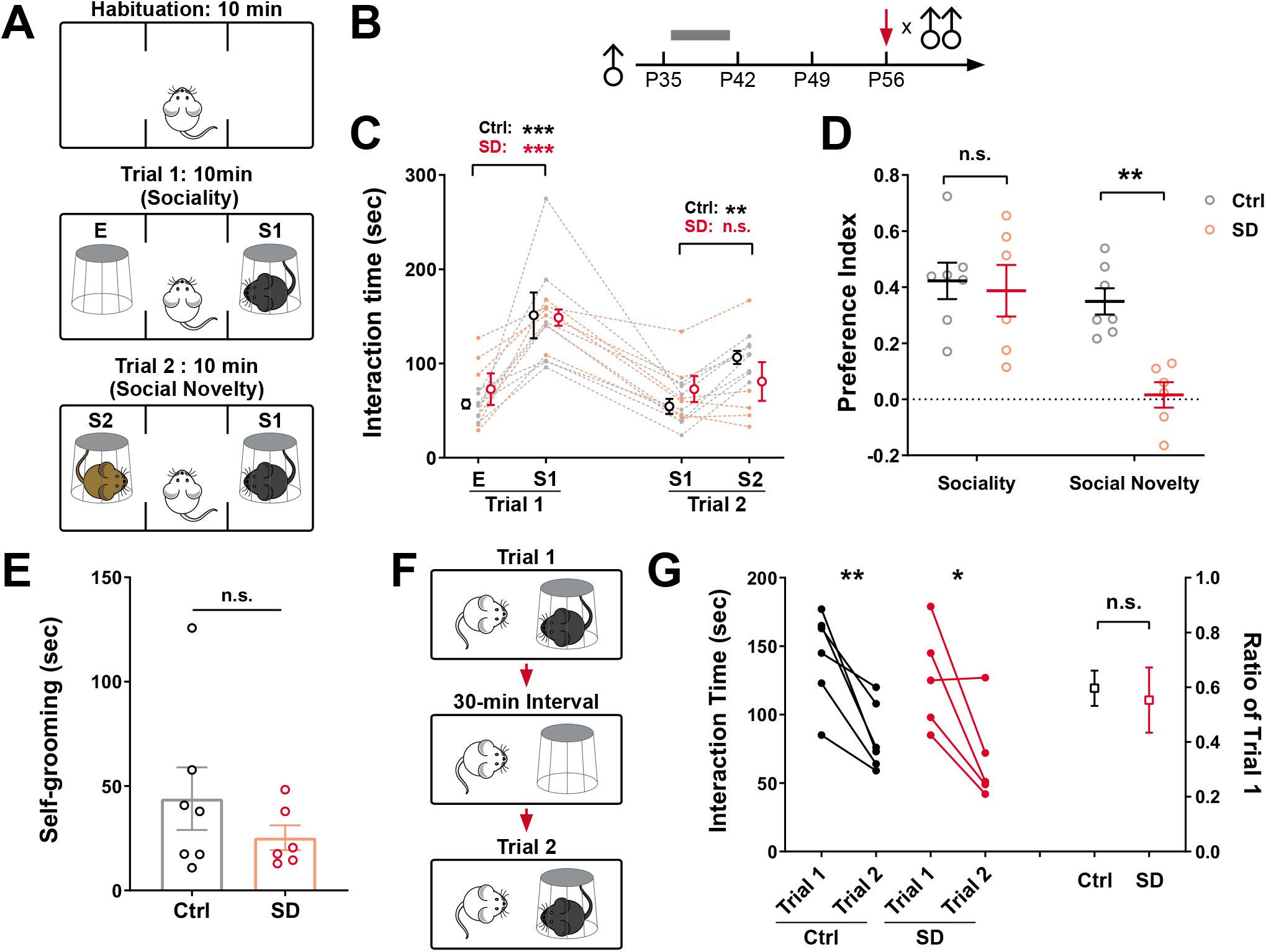
Adolescent SD induced loss of social novelty preference in adult social interactions. (A, B) Diagrams showing the three-chamber social interaction test. (B) Timeline showing that male test mice received daily SD in 5 days during P35 – 42 (grey bar) and were tested for social interactions using male stimulus mice at P56 (red arrow). (C, D) The absolute time (C) and preference indices (D) of interactions with the empty cup (E), stranger 1 (S1) and stranger 2 (S2) in trial 1 and trial 2 (Ctrl, n = 7; SD, n = 6). (E) The time Ctrl or SD mice spent in selfgrooming during the 2 trials of 3-chamber test (Ctrl, n = 7; SD, n = 6). (F) Diagram showing the social memory test. (G) The absolute time of interactions with the stimulus mouse during the social memory test (left) and the ratio of interaction time in trial 2 versus trial 1 (right, Ctrl, n = 6; SD, n = 5). All data are shown as mean ± s.e.m. The *p*-values were calculated using statistical tests listed in Supplementary Table 1. * *p* < 0.05; ** *p* < 0.01; *** *p* < 0.001; n.s., not significant.

The restricted and repetitive behavior (RRB) is another core symptom that often accompanies social defects in autism patients(Association., 2013). Therefore, we also examined a type of RRB in rodents, namely the excessive self-grooming, but did not find a significant difference between the Ctrl and SD mice (Fig. 2C).

The lack of social novelty preference in the SD mice could be due to loss of ability to memorize the social cue from the familiar mouse. To test this possibility, we performed a social memory test during which the test mouse was exposed to the same stimulus mouse but with a 30-minute inter-trial interval (Fig. S2D). However, similar to the Ctrl mice, the SD mice also showed dramatically reduced interaction time during the second trial, suggesting that they can still memorize the stimulus mouse and recognize that it’s not novel anymore after the 30-minute interval (Fig. S2D). Thus, the defect is likely due to lack of preference *per se*, i.e. the motivation to pursue social novelty as rewarding stimuli.

### 3. The impact of adolescent SD is long-lasting, dependent on developmental stages and restricted to same-sex interactions

To determine whether the defect in social novelty preference caused by adolescent SD persisted over time, we re-run the 3-chamber assay using different sets of stimulus mice on the same test mouse 4 weeks after the initial test (P84 – P90, Fig. 3A). At this adult age, Ctrl mice still preferred to interact with the novel stranger rather than the familiar conspecific while the SD mice demonstrated no preference, suggesting the effect of adolescent SD on social novelty preference is long-lasting (Fig. 3A and 3B). We note that at this timepoint, the preference index of sociality in SD mice is lower than that in Ctrl mice despite the SD mice still spent almost 2-fold of time interacting with the stimulus mouse (S1) than the empty cup (E) in Trial 1 (Fig. 3A and 3B). Given that at P56, the sociality preference is almost identical between Ctrl and SD and the preference index is greater than that in SD mice at P84, adolescent SD are likely to accelerate the decay of sociality preference in adulthood, but not its development during adolescence.

**Figure 3.**
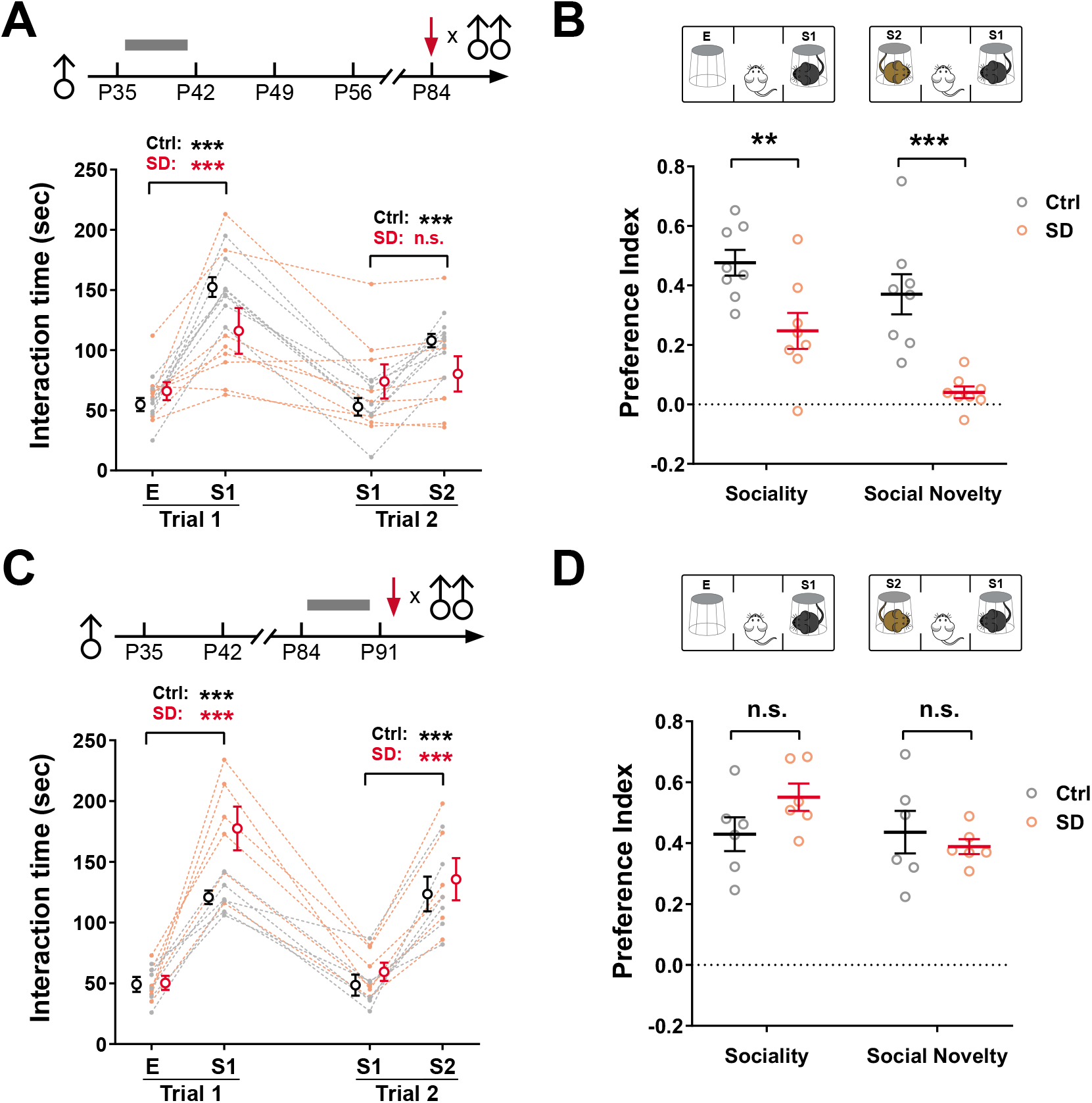
The impact of adolescent SD is long-lasting and development-dependent. (A, B) Male mice receiving Ctrl or SD protocol during P35 – 42, and three-chamber social interaction test was performed using male stimulus mice at ~ P84. Absolute interaction time and preference indices are shown in (A) and (B), respectively (n = 8). (C, D) Male mice receiving Ctrl or SD protocol during P84 – 91, and 3-chamber social interaction test was performed using male stimulus mice at > P91, at least 24 hrs after the last SD session. Absolute interaction time and preference indices are shown in (C) and (D), respectively (n = 6). All data are shown as mean ± s.e.m. The *p*-values were calculated using statistical tests listed in Supplementary Table 1. * *p* < 0.05; ** *p* < 0.01; *** *p* < 0.001; n.s., not significant.

These results raised another interesting question, i.e. can SD impact social interaction behavior regardless whether it occurs during development or not? To address this issue, we applied the same SD and Ctrl protocols in another group of mice when they were already adults (P84 – P91, Fig. 3C) and tested these mice after at least 24 hours of sleep recovery (> P91, Fig. 3C). Interestingly, this adult SD did not cause any defects in either sociality or social novelty preferences (Fig. 3C and 3D), suggesting that undisturbed sleep during adolescence, but not adulthood, is required for shaping the social novelty preference.

Is this effect of adolescent SD specific to male-male interactions? To answer this question, we performed the same adolescent SD protocol on female mice and subsequently probed them for female-female interactions, or on male mice but tested for their interactions with female mice in the adulthood. The SD female mice showed defects in social novelty preference in female-female interactions (Fig. S3A and S3B), similar to what we observed in male-male interactions (Fig. 2A and 2B). However, when encountered with female stimulus mice, the male mice, regardless whether they had previous adolescent SD or not, exhibited strong preference towards the novel female over the familiar one, and the orders of magnitude of changes are comparable between groups (Fig. S3C and S3D). Thus, the novelty preference defect induced by adolescent SD is restricted to same-sex social interactions, and it is likely to be overcome by the instinctive sexual drive during male-female encounters.

### 4. Correlated defects in adolescent sleep and adult social behavior in *Shank3 InsG3680* mice

Transgenic animal models carrying risk mutations identified in human patients of psychiatric disorders often exhibit behavioral abnormalities that recapitulate the symptoms of these disorders. The *Shank3 InsG3680* knock-in (*InsG3680*) mice carry an ASD-associated single guanine insertion at position 3680 of *Shank3* cDNA, causing loss of function of this gene in the mutants. The *InsG3680* mice have been reported having significant defects in both sociality as well as social novelty preference during social interactions (Peca et al., 2011; Zhou et al., 2016). Given the correlation of sleep amount and social communication defects reported in ASD patients (Veatch et al., 2017), we determined whether the *InsG3680* mice also recapitulate these sleep problems, especially during their adolescence and whether the sleep problems correlated with their social performance. We implanted EEG/EMG electrodes in *InsG3680* mice around P28, recorded their spontaneous sleep/wake cycle during adolescence (P35 – P42), and performed 3-chamber social interaction assay in young adulthood (P56 – P60). Consistent with previous studies, homozygous (Homo) mutant mice showed significantly reduced social novelty preference compared to WT littermates (Fig. 4A). However, in our system, the sociality preference seemed to be unaltered in the Homo mice, only showing slight decrease but not significantly different from WT littermates (Fig. 4A). EEG/EMG recordings revealed that Homo mutants showed more waking and less NREM and REM sleep than WT littermates during adolescence (Fig. 4B, 4C, S4A and S4B). These changes of vigilance states occurred mostly in the light phase, with large variation within each group, and therefore only the increase of waking is statistically significant (Fig. 4B, 4C, S4A and S4B). These results indicate abnormal sleep architecture during adolescence in the *InsG3680* mutant mice with large within group heterogeneity.

**Figure 4.**
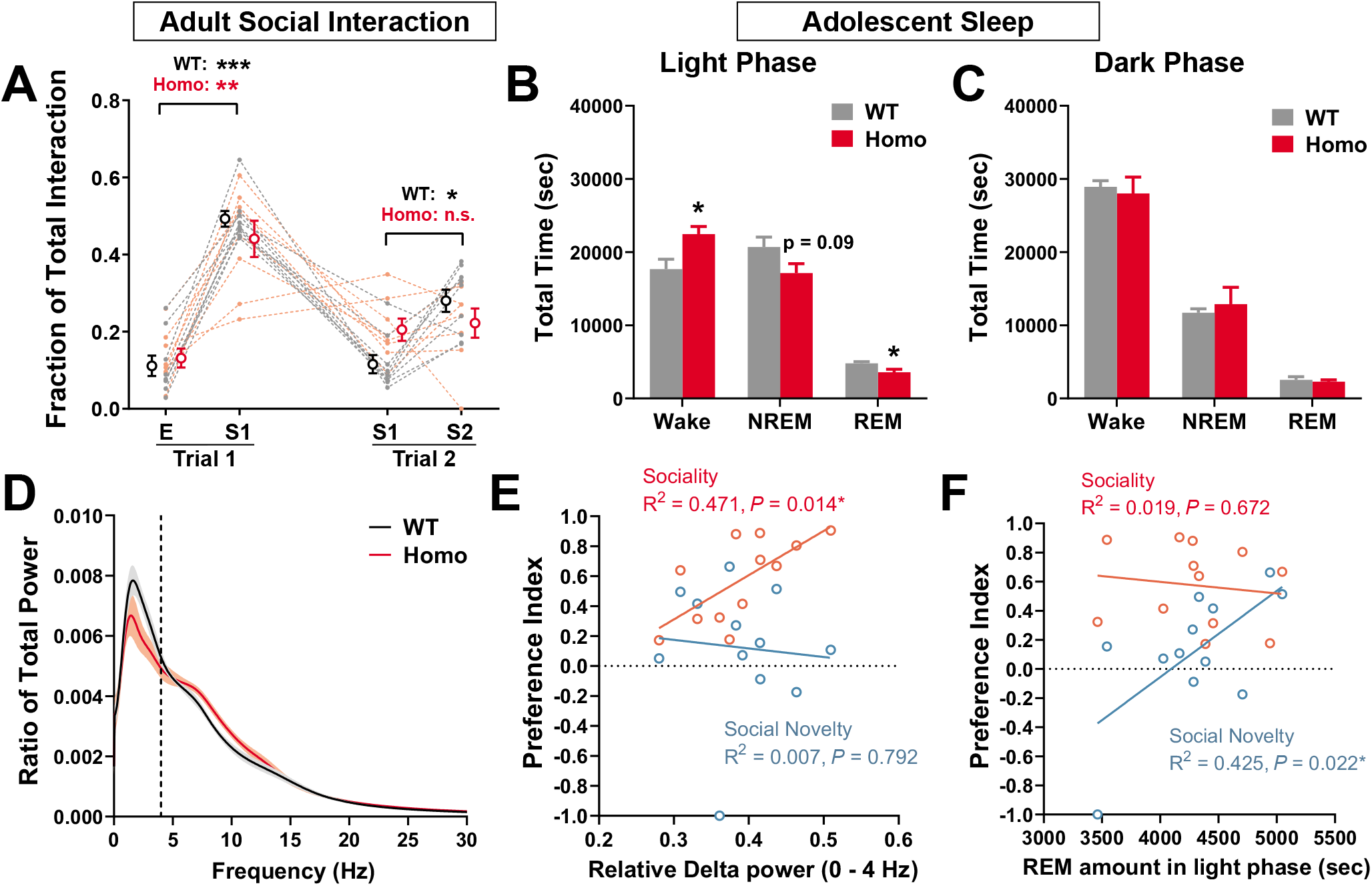
Adolescent sleep defects in *InsG3680* mice correlate to social behavioral deficits in adulthood. (A) Three-chamber social interaction test was performed in homozygous *InsG3680* mice and WT littermates at P56. Interaction time with E, S1 and S2 in each trial was normalized as fraction of total interaction time for each animal (WT, n = 9; Homo, n = 8). (B – D) Total amount of Wake, NREM and REM over 12-hr light phase (B) or dark phase (C) and EEG Power spectrum of NREM (D) at P35 – 42 (WT, n = 5; Homo, n = 6). (E, F) Linear regression between the preference indices of adult social interaction and NREM delta power (E) or light phase REM amount (F) during adolescence. Data points includes 4 WT, 3 Het and 5 Homo mice that had both EEG recording and behavioral tests. All data are shown as mean ± s.e.m. The *p*-values were calculated using statistical tests listed in Supplementary Table 1. * *p* < 0.05; ** *p* < 0.01; *** *p* < 0.001; n.s., not significant.

Given the highly heterogenous sleep phenotypes of *InsG3680* mice, we wondered whether their individual social performance correlated with their adolescent sleep, and more specifically, with a certain aspect of sleep structure. Therefore, we pooled together all sleep and social performance data of the mice that had both EEG/EMG recording and social tests (including 4 WT, 5 Homozygous and 3 Heterozygous mice) and performed linear regression between social preferences and sleep parameters, i.e. amount of Wake, NREM or REM in light phase, and relative Delta (0 – 4 Hz), Theta (4 – 7 Hz), Alpha (7 – 12 Hz) or Beta (12 – 30 Hz) power of NREM sleep (Fig. 4E, 4F and S4C-S4G). We found that the relative Delta power of NREM shows a strong positive correlation with sociality preference but not with social novelty preference (Fig. 4E). However, relative Alpha power and Beta power of NREM are negatively correlated with sociality preference (Fig. S4E-S4G), but also not with social novelty preference, although these wavebands only take small fractions in NREM EEG. Interestingly, on the other hand, the amount of REM sleep in light phase shows a positive correlation with social novelty preference but not with sociality preference (Fig. 4F), while neither of Wake nor NREM amount shows correlation with the social performance (Fig. S4C and S4D). Together, these results demonstrate specific association of a particular sleep component in adolescence with a distinct aspect of adult social behavior, and further suggest that REM sleep may be more important for the shaping of social novelty preference while NREM sleep is more linked to sociality preference.

## Discussion

### A critical period for shaping the neural network underlying social behaviors

The functions of sleep during early postnatal phase (i.e. childhood) and later developmental phase (i.e. adolescence) are likely to be different from each other and distinct from that in adulthood. However, studying sleep in early development is extremely difficult because of technical limitations, especially when one wants to causally associate it with behavioral outcomes in adulthood. Adolescence, on the other hand, provides an ideal time window that sleep monitoring and manipulations are feasible and the adolescent brain still undergoes substantial developmental changes such as synapse remodeling and circuit refinement(Bhatt et al., 2009; Bian et al., 2015; Moyer and Zuo, 2018). A study using chronic sleep restriction during the early postnatal stage in mice (P5 – P42) showed that it induced a series of long-lasting behavioral changes 4 weeks after the completion of sleep restriction, including hypoactvity in both sexes, slightly increased sociality and social novelty preference in female mice but not in male mice, and female-specific decrease of marble burying behavior(Sare et al., 2016). However, a most recent follow-up study from the same lab showed that the same postnatal sleep restriction in male mice indeed led to changes of social interaction behaviors, with sociality increased but social novelty preference impaired(Sare et al., 2019). The sleep restriction method used in these studies could be potentially problematic because its effects on sleep cannot be validated by EEG in mouse pups and the influence of maternal separation at this young age cannot be excluded, however results from these studies somewhat echoes with ours showing impairment only in social novelty preference but not in sociality preference by SD at later adolescence (Fig. 2A and 2B), suggesting that distinct neural circuits underlie the shaping of these behavioral preferences and are vulnerable to sleep interventions at distinct developmental stages. More interestingly, both sleep manipulations, regardless in early childhood(Sare et al., 2016; Sare et al., 2019) or during adolescence (Fig. 3A and 3B), resulted to long-lasting behavioral changes in social interactions, highlighting the idea that sleep helps to shape the neural network underlying social behavior when it is still plastic during development, and the effects become “fixed” as the neural network matures in adulthood.

This is further supported by our finding that after a certain age point (> P56 in this study) sleep disruption no longer has the ability to influence social interaction behaviors (Fig. 3C and 3D). This development-dependent effect is reminiscent of the “critical period” plasticity in the developing visual system(Hensch, 2005), during which the outer stimuli or experiences impact the functional outcome to a much greater extent compared to the same stimuli/experiences before or after this period. The critical period plasticity has been found in numerous biological processes including the separation of olfactory glomeruli, refinement of whisker responsive fields in the somatosensory cortex, remodeling of synapses at retina, visual thalamus and cortex, as well as acquisition of motor skills(Fox and Wong, 2005; Peters et al., 2017; Petersen, 2007; Yang et al., 2018). Therefore, we argue that the shaping of social behavior, especially the social novelty preference and the underlying neural circuits also possesses a “critical period” which is within adolescence.

From a canonical view, a specific neural network carrying a certain function can be refined by experiences/stimuli of the same modality, e.g., visual experience for visual circuits, motor training for motor circuits, social experiences for social circuits (Bian et al., 2015; Feldman and Brecht, 2005; Fox and Wong, 2005; Fu et al., 2012; Li et al., 2017b; Remedios et al., 2017; van der Bourg et al., 2019; Xu et al., 2009). However, our study demonstrates a different type of neural network modification or refinement which is induced by global activity pattern or brain state changes. Whether alteration of adolescent sleep impact other aspects of brain functions, e.g. sensory processing, motor functions, or learning and memory, and though what mechanisms remain open questions and call for further investigation.

### Links between specific sleep components during adolescence and social behaviors in adulthood

Here we show in the *Shank3 InsG3680* knock-in strain, the Homo mice have defects in both adolescent sleep and adult social interactions, despite large variations in both phenotypes within each genotype(Fig. 4A-D, and S4A, B). We further demonstrate the correlations between specific sleep components and the social preferences, i.e. significant positive correlations between NREM delta power and sociality preference, and between light phase REM sleep amount and social novelty preference (Fig. 4E and 4F). Since REM sleep mostly occurs during the light phase and the REM amount was almost not changed in the dark phase, the social novelty preference should also correlate with total REM amount across the 24-hour cycle. Interestingly, the sociality preference is also negatively correlated with powers of higher frequency bands in NREM, i.e. Alpha (7 – 12 Hz) and Beta (12 – 30 Hz) (Fig. S4F and S4G). Thus, all parameters that correlate with sociality preference are NREM components while social novelty preference is only correlated with REM sleep. In WT C57BL6 mice, although both NREM and REM sleep are deprived by our adolescent SD protocol, we achieved 100 % deprivation of REM sleep with relatively weak rebound after the SD session whereas ~15% of NREM sleep was still retained in the last 2 hours of SD and had more dramatic rebound afterwards. Therefore, the social novelty preference may be more sensitive to REM sleep depletion, than NREM sleep reduction, in adolescence.

Although the 12 – 30 Hz band only takes a small fraction of NREM power spectrum, it overlaps with the frequency range of sleep spindles (usually 12 – 14 Hz) which, together with the sharp wave/ripple activity in the hippocampus and cortical slow waves, have been linked with memory consolidation and cognitive performance(Diekelmann and Born, 2010; Langille, 2019; Latchoumane et al., 2017). However, in our results the 12 – 30 Hz band power showed correlation with sociality opposite to that of delta band which contains the slow wave activity (0.5 – 4 Hz), suggesting these sleep activities play different roles in shaping the social preference than that in learning and memory.

## Acknowledgements

We acknowledge all de Lecea lab members for critical feedback. We thank A. Khan for technical assistance. This work was supported by Human Frontier Science Program fellowship LT000338/2017-L (W.-J.B.) and National Institutes of Health grants R01 MH102638 (L.d.L.), R01 MH087592 (L.d.L.) and R01 MH116470 (L.d.L.). We also acknowledge support from A. Olson and the Stanford Neuroscience Microscopy Service, NIH NS069375.

## Author contributions

W.-J.B. and L.d.L. designed the project. W.-J.B. performed experiments and data analyses. W.-J.B. wrote the manuscript with contributions from L.d.L.

## Experimental Procedures

### Animals

All experimental protocols were approved by the Stanford University Animal Care and Use Committee and in accordance with the National Institutes of *Health Guide for the Care and Use of Laboratory Animals*. The *Shank3 InsG3680* knock-in mice (*InsG3680*, full name: STOCK *Shank3^tm3.1Gfng^*/J; JAX strain 028778; gift of Prof. Guoping Feng, Massachusetts Institute of Technology, U.S.A.)(Zhou et al., 2016) were kept on 129S2/SvPasCrl background. Offspring of *Shank3 InsG3680* mice or C57BL6J wildtype breeders were born and housed at constant temperature (22 ± 1 °C) and humidity (40–60%), under a 12/12-hour light–dark cycle (lights-on: 7:00 a.m. – 19:00 p.m., ZT 0 – 12; lights-off: 19:00 p.m. – 7:00 a.m., ZT 12 – 24). The mouse pups were weaned on postnatal day 21 (P21) and subject to experimental procedures after they reached adolescence (⩾ P28). Mice were allowed to access food and water *ad libitum*.

### Developmental Sleep Disruption

The sleep disruption (SD) was achieved using the automatic Sleep Deprivation System (ViewPoint Life Sciences, Inc., Lyon, France) which is composed of a deprivation chamber (PVC cylinder, Height: 46 cm, Width: 30 cm, Weight: 5 Kg) with a shaking platform at the bottom, a controller and a laptop with the controlling program installed. Mice were transferred to the deprivation chamber where they had access to food and water *ad libitum* and returned to the homecage after daily SD or Ctrl sessions. Programed electromagnetic pulses were delivered to the shaking platform and controlled its movement to keep the mice awake during the sessions. The parameters of electromagnetic pulses were as follow: SD, randomized 2 – 8 pulses (15-ms duration) delivered at 2 Hz every 0.2 – 0.8 minutes, for 4 hours daily during early light phase (ZT 2 – 6); Ctrl, 5 pulses of the same duration delivered at 2 Hz every 0.5 minutes, for 4 hours daily during early dark phase (ZT 12 – 16). We avoided performing the SD protocol in the very beginning of light phase (ZT 0 – 2) in order not to induce drift in circadian rhythm. For mice with simultaneous EEG/EMG recording, a divider was placed in the deprivation chamber to separate the mice from each other, and up to 4 mice were sleep-deprived/recorded at the same time. The parameters of SD protocol were deliberately tuned not to induce acute stress, as confirmed by ELISA of plasma corticosterone level after the SD session (Fig. 1J). The SD or Ctrl protocol were performed for 5 consecutive days between P35 and P42, then the mice were left undisturbed for 2 weeks and the behavioral tests were carried out after P56.

For the “gentle-touch” procedure, mice stayed in their homecage and were monitored by an experimenter (W.-J. B.). If an animal remains motionless for a few seconds, he was gently touched with a soft brush. Novel objects (paper tubes, cotton nestlets, etc) were also added into the homecage to help keeping the mice awake but removed after the SD session. As control, same numbers of touching and objects were given to the Ctrl mice during the dark phase.

For measurement of plasma corticosterone, mice were anesthetized using isoflurane and a small quantity of blood sample (~ 100 μl) were collected from the orbital sinus on the first and last day of SD and immediately after the 4-hour SD session. Control samples were collected from naïve animals without any manipulation. Plasma were separated from the whole blood sample by centrifuge and proceeded to ELISA according to manufacture’s instructions (Enzo, ADI-900-097).

### Surgery

Surgery for EEG/EMG implantation was carried out at P28 – P30. The animal received a subcutaneous injection of Buprenorphine SR (1mg/kg) before incision and was anesthetized with a mix of ketamine (100mg/kg) and xylazine (10 mg/kg) injected intraperitoneally (i.p.). The animal was then placed on a stereotaxic rig (David Kopf Instruments, Tujunga, CA). Cortical EEG and EMG electrodes were implanted as described in our previously studies(Eban-Rothschild et al., 2016; Li et al., 2018). Briefly, stainless steel mini-screws (US Micro Screw) for EEG were implanted to the skull above the frontal (AP–1.6 mm; ML 1 mm) and temporal (AP 3 mm; ML 2.5 mm) lobes. Mini-rings made of metal wires (316SS/44T, Medwire) were inserted into neck muscles for EMG recording. The electrodes were previously soldered to a 4-pin connector which was mounted on the skull using Metabond (Parkell) and dental cement.

### EEG/EMG recording and data analysis

After the surgery, the animal was allowed to recover for at least 1 week and connected to a flexible recording cable at least 24 hours before the recording began. EEG and EMG recording across a complete 24-hour light-dark cycle was performed on the day before SD started (Baseline), on a single day within the 5-day SD period (During SD), or 24 hours after the last SD day (After SD). For *InsG3680* mice, EEG/EMG recordings were carried out between P35 – P42. EEG/EMG signals were amplified through a multi-channel amplifier (Grass Instruments) and collected by VitalRecorder (Kissei Comtec Co.) at sampling rate of 256 Hz filtered between 0 and 120 Hz for offline signal analysis. Raw EEG/EMG data were converted in SleepSign (Kissei Comtec Co.), exported to Matlab (MathWorks) and analyzed with custom Matlab scripts(Li et al., 2018). Power-frequency distributions between 0.5 – 30 Hz frequency range of the NREM episodes were analyzed using the power spectral density (PSD) function in the Matlab scripts, and the relative power of delta (0.5 – 4 Hz), theta (4 – 7 Hz), alpha (7 – 12 Hz) and beta (12 – 30 Hz) bands was calculated as fraction of total power (0.5 – 30 Hz) for each individual animal.

### Behavioral assays

All behavioral assays were carried out in early dark phase (ZT 12 – 16) in a dark experiment room with only dim red lights to minimize the acute disruption to sleep and circadian rhythm, except for the novel object recognition test which requires visual cues to help to discriminate the objects and therefore was performed in the last 2 hours of light phase (ZT 10 – 12) with lights on. Mice were habituated to the experiment room at least 1 hour before the test started. *Three-chamber social interaction Test*. The experimental procedure was as described elsewhere (Kaidanovich-Beilin et al., 2011). In brief, as shown in Fig 2A and S1A, the test was performed in an apparatus of 3 compartment chambers (2 side chambers: 26 cm × 23 cm; middle chamber: 11 cm × 23 cm) with connecting doors. For all experiments, the test was performed in early dark phase (ZT12 – 16). For habituation, the test mouse was placed in the middle chamber and allowed 10 minutes of free exploration of all 3 chambers in the empty apparatus. The mouse was then returned to the middle chamber and doors covered by cardboards. An empty metal mesh cup (Empty, E) was placed randomly in one of the 2 side chambers, serving as the non-social novel object, and another identical mesh cup containing a never-before-met stimulus mouse (Stranger 1, S1) was placed in the opposite chamber. The covering of doors was then removed, and the test mouse was free to explore and interact with either E or S1 for 10 minutes (Trial 1). After Trial 1 was completed, a second never-before-met mouse (Stranger 2, S2) was placed in the previously empty cup, serving as the social novelty stimulus while the S1 had become familiar. The test mouse was again allowed to explore both stimulus mice for another 10 minutes (Trial 2). Trial 1 and Trial 2 were videotaped for analysis. The time spent in direct interactions (e.g. sniffing, touching with head and forelimbs) between the test mouse with either E, S1 and S2 in each trial was measured manually using a stopwatch. For *InsG3680* mice, due to the large within-group variation, the interaction time with E, S1 or S2 was normalized using the total time spent interacting with either of them during the entire test session (20 minutes) for each animal. Preference index of sociality or social novelty (SN) was calculated as follows, *PI_Sociality_* = *(t_S1_ – t_E_)/(t_S1_ + t_E_); PI_SN_ = (t_S2_ – t_S1_)/(t_S2_ + t_S1_)*, where *t_E_, t_S1_* and *t_S2_* are the time of interaction with E, S1 and S2, respectively, within the given trial. Total number of door-crossing and the time spent in self-grooming of the test mouse was also quantified for the whole test session (Trial 1 + Trial 2, 20 minutes in total). Stimulus mice of same sex with the test mice were used for all social interaction experiments except for examining male-female interaction, where the male mice were used for test subjects and female mice were used for stimulus objects.

#### Novel object recognition (NOR) with reduced memory component

NOR assay typically includes a training trial, a delay period of 24 hours for memory consolidation, and a test trial(Li et al., 2018; Rolls et al., 2011). However, in order to better examine the preference *per se*, we conducted this assay using a protocol with reduced memory component (Fig. S2E). After 10 minutes of habituation to the apparatus (black walled open arena, 54 cm × 26 cm), mice were given two identical non-social objects for exploration for 10 minutes (Training). Each object was placed at the same distance from the walls and corners of the arena with no specific spatial or odor cues. Immediately after this 10-minute training trial, one of the objects was replaced with a novel, non-social object (Test), and the animal was allowed another 5 minutes of exploration. Training and Test trials were videotaped for analysis, and the direct interaction (e.g. sniffing, touching with head and forelimbs) between the mouse and each object during the first 5 minutes of each trial were quantified. Mice that demonstrated clear bias during the Training trial (> 65% preference for either object) were excluded from the experiment.

#### Social memory test

The experimental procedure was as described elsewhere(Winslow, 2003). The test mouse was placed in an open arena (same with that used in NOR) containing an empty mesh cup (identical to that used in the three-chamber test) for habituation of 10 minutes. A novel, stimulus mouse of same sex was then placed into the mesh cup, and the test mouse was allowed to explore the stimulus mouse and their interactions were videotaped for 5 minutes (Trial 1). The stimulus mouse was then removed, and the test mouse was left alone in the arena for 30 minutes. After this 30-minute interval, the same stimulus mouse was put back in the mesh cup, and the interactions will be recorded for another 5 minutes (Trial 2). Similar to the three-chamber test, the time of direct social interaction was quantified for each trial.

#### Elevated plus maze (EPM)

EPM is a well-established assay measuring anxiety in laboratory rodents(Walf and Frye, 2007). It was done in an elevated maze with four arms (two open and the other two closed, each arm is 66 cm long and 5 cm wide). Mice were placed at the junction of the open and closed arms, facing the open arm opposite to the experimenter. Mice were allowed to explore the maze for 5 minutes. Due to occurrence of falling from the open arms during our experiments (3 out of 8 in SD group), we quantified the latency to first entry into the open arms instead of total time spent in the open arms.

### Statistical analysis

Statistical tests were carried out using GraphPad Prism 8 (GraphPad software). Two-tailed Welch’s t-tests were used for comparison between 2 conditions, except for Fig 1D and 1G, where paired t-tests were used. One-way ANOVA followed by Dunnett’s or Tukey’s multiple comparison tests were used for comparison of 3 or more conditions. For experiments with two independent variables, two-way ANOVA followed by Bonferroni’s or Tukey’s post-tests were used, depending on the numbers of conditions of the post-tests. Repeated measures were incorporated when appropriate (RM-ANOVA). The simple linear regression function in GraphPad Prism 8 (GraphPad software) was used to evaluate the correlation between sleep components and social preferences in *InsG3680* mice. Only data from the animals that had both sleep recordings and behavioral tests were included in this analysis. Sleep states or EEG wavebands are considered categories but not independent variables (Fig. 1D–1I, 2B and 2C), therefore t-tests or one-way ANOVA were used for comparisons within each state or waveband. A detailed list of statistical tests used for each figure panel, including exact F and *p* values, was in Supplementary Table 1. Data distribution was assumed to be normal although not formally tested. All data are presented as mean ± s.e.m. Sample sizes were predetermined using webbased sample size/power calculator (https://www.stat.ubc.ca/~rollin/stats/ssize/n2.html) and are similar to those reported in previous publications in the field (Cao et al., 2018; Eban-Rothschild et al., 2016; Giardino et al., 2018; Hung et al., 2017; Li et al., 2018; Sgritta et al., 2019). All behavioral analyses were performed blinded to the experimental conditions. All conditions statistically different from control are indicated. * P < 0.05; ** P < 0.01; *** P < 0.001.

