## Supplementary Figures and Legends for "Adolescent sleep is critical for the developmental shaping of social novelty preference"

**Supplementary Figure 1. Developmental shaping of social preferences.**

(A) Diagrams showing the 3-chamber social interaction test. (B, C) Naïve WT C57B6 mice of different age shows development-dependent increases of sociality preference (B) and social novelty preference (C) during the 3-chamber social interaction tests. Preference indices were calculated as indicated in each graph (n = 4). All data are shown as mean ± s.e.m.

**Supplementary Figure 2. Behavioral probing of adolescent SD mice.**

(A) Data points of individual mice that received different SD methods between P35 – 42 are plotted in different color. Same animals with Fig. 2C, D and E (n = 6). (B) Male mice received Ctrl or SD between P42 – 49 and were assayed at P56 by the three-chamber test using male stimulus mice (Ctrl, n = 4; SD, n = 5, RM two-way ANOVA followed by Tukey's multiple comparisons test). (C) 10 – 12-week-old mice receiving SD at P35 – 42 showed similar latency to their first entry to the open arm when placed in an elevated plus maze (Ctrl, n = 9; SD, n = 8, unpaired t test with Welch's correction). The 2 mice showing latency of 300 sec did not enter the open arm at al during the whole 5-min assay. (D) Mice receiving previous Ctrl or SD at P35 – 42 showed similar total number of door-crossing during the whole 3-chamber social interaction test (Ctrl, n = 7; SD, n = 6, unpaired t test with Welch's correction. Same animals with Fig. 2C, D and E). (E) Diagram of the novel object recognition assay. (F) The interaction time with the familiar and the novel objects during the Test session of novel object recognition assay (Ctrl, n = 6; SD, n = 5, RM two-way ANOVA followed by Bonferroni's multiple comparisons test, Same animals with Fig. 2G). All data are shown as mean ± s.e.m. * $p < 0.05$; ** $p < 0.01$; *** $p < 0.001$; n.s., not significant.

**Supplementary Figure 3. The social interaction defect induced by adolescent SD is restricted to same-sex interactions.**

(A, B) Female mice received Ctrl or SD between P35 – 42 and were assayed at P56 by 3-chamber test using female stimulus mice (n = 6). (C, D) Male mice received Ctrl or SD between P35 – 42 and were assayed at P56 by 3-chamber test using OVXed female stimulus mice (n = 6). Interaction time (RM two-way ANOVA followed by Tukey's multiple comparisons test) and preference indices of sociality and social novelty (RM two-way ANOVA followed by

Bonferroni's multiple comparisons test) are shown in (A, C) and (B, D), respectively. All data are shown as mean ± s.e.m. * $p < 0.05$; ** $p < 0.01$; *** $p < 0.001$; n.s., not significant.

**Supplementary Figure 4. The adolescent sleep components are associated with adult social preferences in *Shank3 InsG3680* mice.**

(A, B) Percentages of time spent in NREM and REM sleep over the 24-hr light-dark cycle in *InsG3680* mice (WT, n = 5; Homo, n = 6). (C - G) Linear regressions of light phase Wake amount (C), light phase NREM amount (D) , NREM theta (E), alpha (F) and beta (G) power with sociality and social novelty preferences ($n = 12$ including 4 WT, 3 Het and 5 Homo). All data are shown as mean ± s.e.m. * $p < 0.05$; ** $p < 0.01$; *** $p < 0.001$; n.s., not significant.

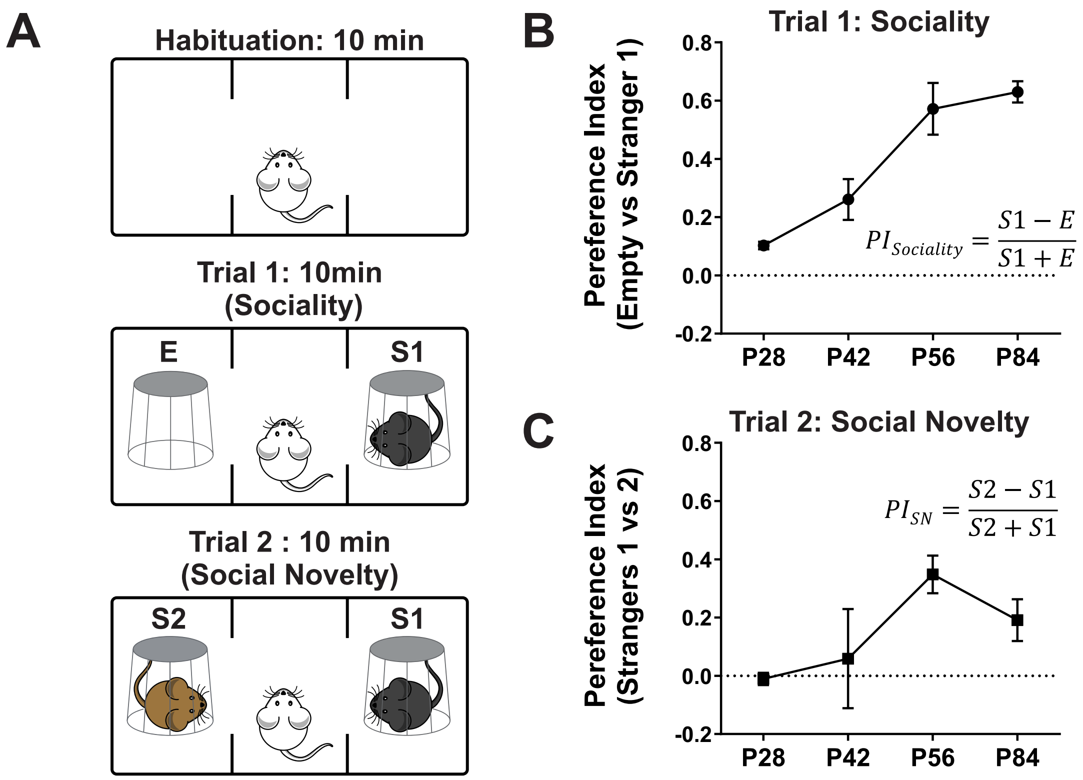


Supplementary Figure 1. Bian et al.

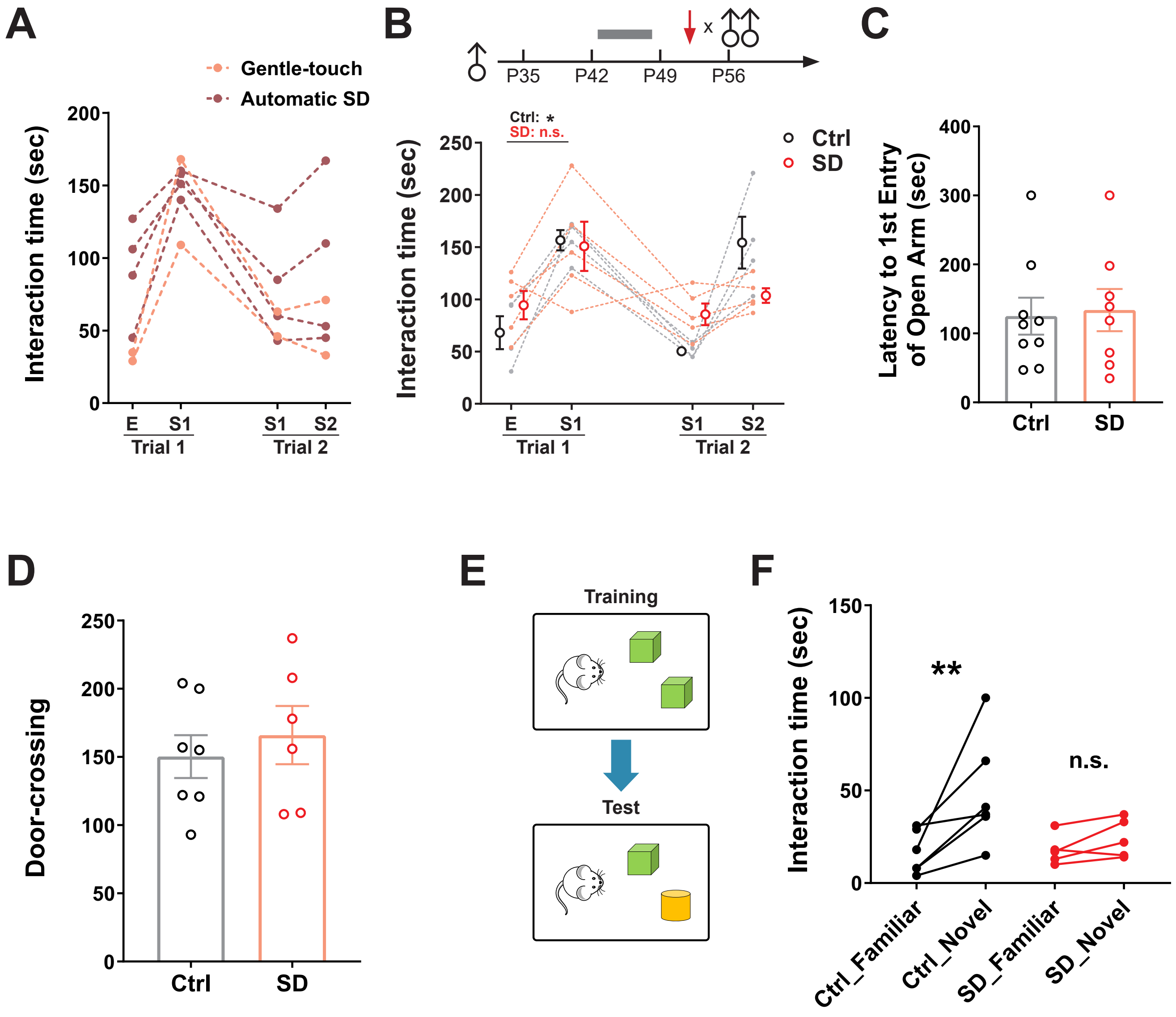


Supplementary Figure 2. Bian et al.

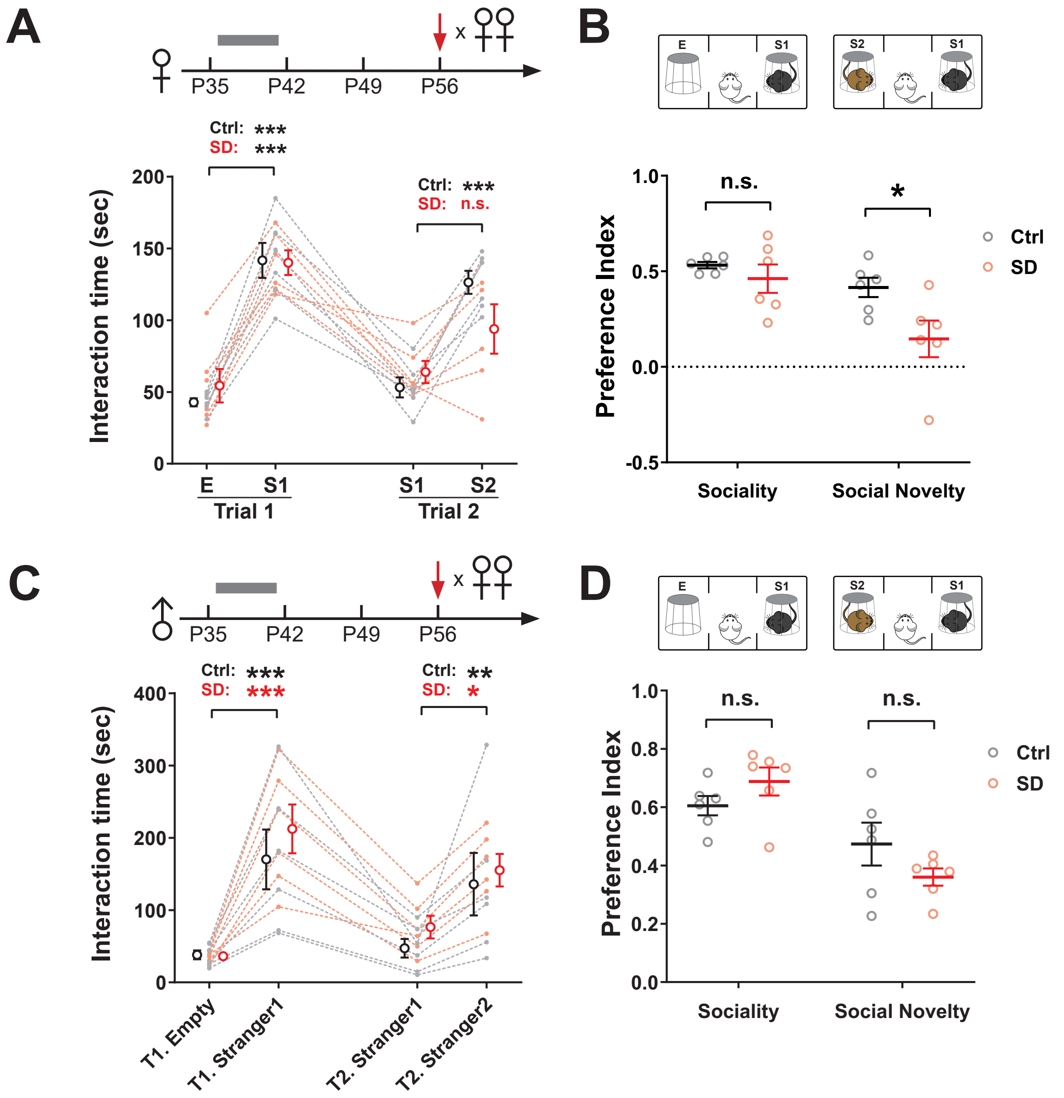


Supplementary Figure 3. Bian et al.

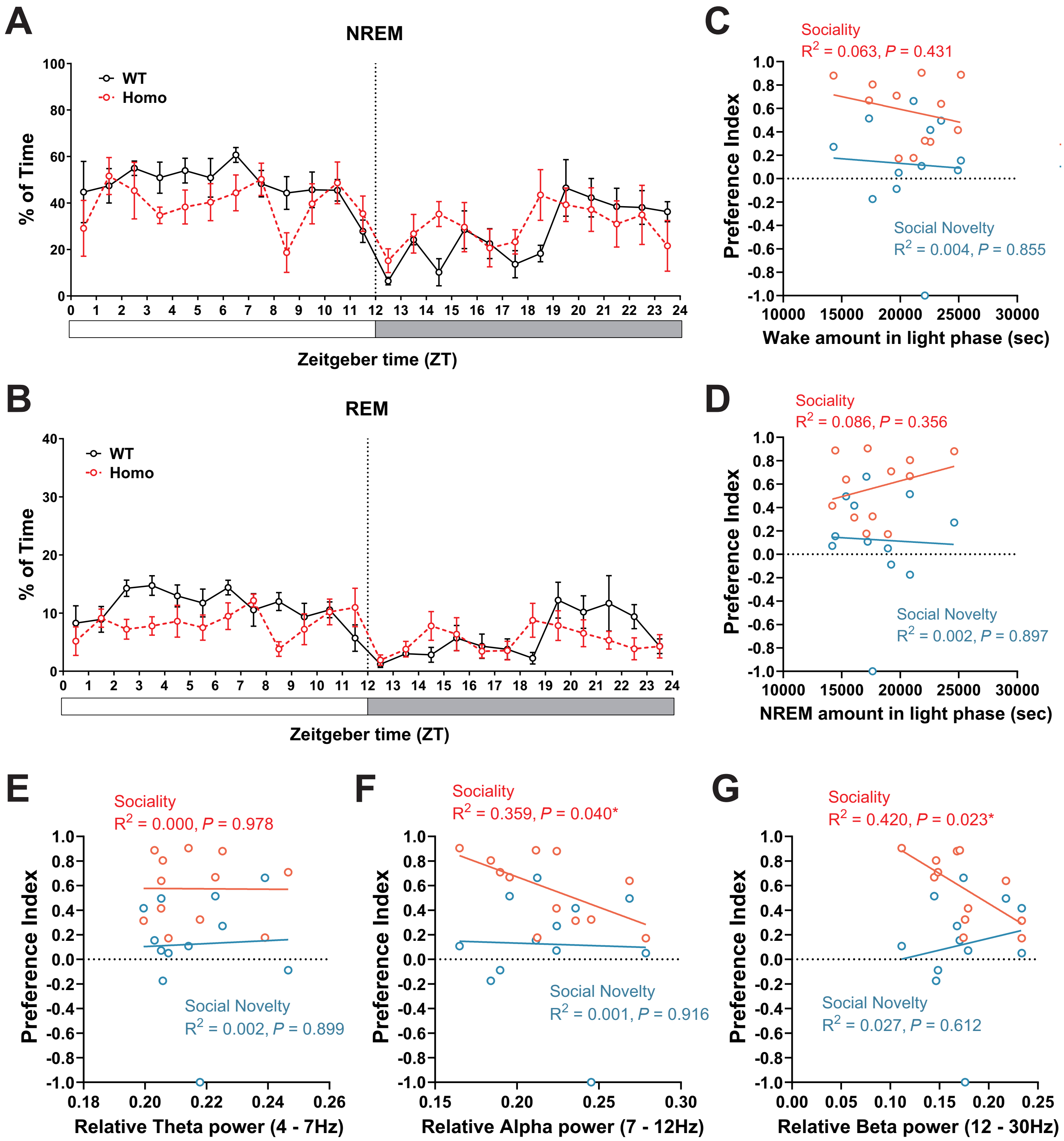


Supplementary Figure 4. Bian et al.
