## Supplementary Table 1 for "Adolescent sleep is critical for the developmental shaping of social novelty preference"

| Figure | Sample size | Statistic test | P value | | |
|---|---|---|---|---|---|
| 1A | 4 | Two-way RM ANOVA | Time x Treatment, F (46, 207) = 2.895 | P<0.0001 | |
| | | Factor 1: Treatment | Time, F (23, 207) = 11.71 | P<0.0001 | |
| | | (Baseline, During SD, | Treatment, F (2, 9) = 27.60 | P=0.0001 | |
| | | After SD) | Multiple comparisons P: | | |
| | | Factor 2: Time | | Baseline vs. During SD | Baseline vs After SD |
| | | Bonferroni's multiple | ZT0-1 | >0.9999 | >0.9999 |
| | | comparisons test | ZT1-2 | >0.9999 | >0.9999 |
| | | | ZT2-3 | 0.0102 | >0.9999 |
| | | | ZT3-4 | 0.0007 | >0.9999 |
| | | | ZT4-5 | 0.0074 | >0.9999 |
| | | | ZT5-6 | 0.0306 | >0.9999 |
| | | | ZT6-7 | >0.9999 | >0.9999 |
| | | | ZT7-8 | >0.9999 | >0.9999 |
| | | | ZT8-9 | >0.9999 | >0.9999 |
| | | | ZT9-10 | >0.9999 | >0.9999 |
| | | | ZT10-11 | >0.9999 | >0.9999 |
| | | | ZT11-12 | >0.9999 | >0.9999 |
| | | | ZT12-13 | >0.9999 | >0.9999 |
| | | | ZT13-14 | >0.9999 | >0.9999 |
| | | | ZT14-15 | >0.9999 | >0.9999 |
| | | | ZT15-16 | >0.9999 | 0.024 |
| | | | ZT16-17 | >0.9999 | >0.9999 |
| | | | ZT17-18 | >0.9999 | >0.9999 |
| | | | ZT18-19 | >0.9999 | >0.9999 |
| | | | ZT19-20 | >0.9999 | >0.9999 |
| | | | ZT20-21 | 0.45 | >0.9999 |
| | | | ZT21-22 | >0.9999 | >0.9999 |
| | | | ZT22-23 | >0.9999 | >0.9999 |
| | | | ZT23-24 | >0.9999 | >0.9999 |
| 1B | 4 | Two-way RM ANOVA | Time x Treatment, F (46, 207) = 3.227 | P<0.0001 | |
| | | Factor 1: Treatment | Time, F (23, 207) = 12.38 | P<0.0001 | |
| | | (Baseline, During SD, | Treatment, F (2, 9) = 3.345 | P=0.0820 | |
| | | After SD) | Multiple comparisons P: | | |
| | | Factor 2: Time | | Baseline vs. During SD | Baseline vs After SD |
| | | Bonferroni's multiple | ZT0-1 | >0.9999 | >0.9999 |
| | | comparisons test | ZT1-2 | >0.9999 | >0.9999 |
| | | | ZT2-3 | 0.0008 | >0.9999 |
| | | | ZT3-4 | 0.002 | >0.9999 |
| | | | ZT4-5 | 0.0004 | >0.9999 |
| | | | ZT5-6 | 0.0017 | 0.631 |
| | | | ZT6-7 | >0.9999 | >0.9999 |
| | | | ZT7-8 | >0.9999 | >0.9999 |
| | | | ZT8-9 | >0.9999 | >0.9999 |
| | | | ZT9-10 | >0.9999 | >0.9999 |
| | | | ZT10-11 | >0.9999 | >0.9999 |
| | | | ZT11-12 | >0.9999 | >0.9999 |
| | | | ZT12-13 | >0.9999 | >0.9999 |

| | | | |
|---|---|---|---|
| | | | ZT13-14 >0.9999 >0.9999<br>ZT14-15 >0.9999 >0.9999<br>ZT15-16 >0.9999 >0.9999<br>ZT16-17 >0.9999 >0.9999<br>ZT17-18 >0.9999 >0.9999<br>ZT18-19 >0.9999 >0.9999<br>ZT19-20 >0.9999 >0.9999<br>ZT20-21 >0.9999 >0.9999<br>ZT21-22 >0.9999 >0.9999<br>ZT22-23 >0.9999 >0.9999<br>ZT23-24 >0.9999 >0.9999 |
| 1D | 9 | Paired t-test | Delta (0 - 4Hz) P = 0.5249<br>Theta (4 - 7Hz) P = 0.2945<br>Alpha (7 - 12Hz) P = 0.5567<br>Beta (12 - 30Hz) P = 0.9734 |
| 1E_Wake | 9 | One-way RM ANOVA<br>Dunnett's multiple comparisons test | Treatment, F (1.718, 13.75) = 54.78 P<0.0001<br>Multiple comparisons:<br>Baseline vs. During SD P<0.0001<br>Baseline vs. After SD P = 0.9342 |
| 1E_NREM | 9 | One-way RM ANOVA<br>Dunnett's multiple comparisons test | Treatment, F (1.868, 14.94) = 70.18 P<0.0001<br>Multiple comparisons:<br>Baseline vs. During SD P<0.0001<br>Baseline vs. After SD P = 0.9224 |
| 1E_REM | 9 | One-way RM ANOVA<br>Dunnett's multiple comparisons test | Treatment, F (1.423, 11.38) = 58.42 P<0.0001<br>Multiple comparisons:<br>Baseline vs. During SD P<0.0001<br>Baseline vs. After SD P = 0.2298 |
| 1F_Wake | 9 | One-way RM ANOVA<br>Dunnett's multiple comparisons test | Treatment, F (1.479, 11.83) = 230.2 P<0.0001<br>Multiple comparisons:<br>Baseline vs. During SD P<0.0001<br>Baseline vs. After SD P = 0.9460 |
| 1F_NREM | 9 | One-way RM ANOVA<br>Dunnett's multiple comparisons test | Treatment, F (1.279, 10.24) = 167.7 P<0.0001<br>Multiple comparisons:<br>Baseline vs. During SD P<0.0001<br>Baseline vs. After SD P = 0.9950 |
| 1F_REM | 9 | One-way RM ANOVA<br>Dunnett's multiple comparisons test | Treatment, F (1.776, 14.21) = 79.99 P<0.0001<br>Multiple comparisons:<br>Baseline vs. During SD P<0.0001<br>Baseline vs. After SD P = 0.0977 |
| 1G_Wake | 9 | One-way RM ANOVA<br>Dunnett's multiple comparisons test | Treatment, F (1.008, 8.065) = 24.47 P=0.0011<br>Multiple comparisons:<br>Baseline vs. During SD P = 0.0018<br>Baseline vs. After SD P = 0.9769 |
| 1G_NREM | 9 | One-way RM ANOVA<br>Dunnett's multiple comparisons test | Treatment, F (1.397, 11.18) = 67.54 P<0.0001<br>Multiple comparisons:<br>Baseline vs. During SD P<0.0001<br>Baseline vs. After SD P = 0.9814 |
| 1G_REM | 9 | Paired t-test | Baseline vs. After SD P = 0.3366 |

| | | | |
|---|---|---|---|
| 1H_Wake | 9 | One-way RM ANOVA<br>Dunnett's multiple comparisons test | Treatment, F (1.601, 12.81) = 11.30 P=0.0023<br>Multiple comparisons:<br>Baseline vs. During SD P = 0.0021<br>Baseline vs. After SD P = 0.9990 |
| 1H_NREM | 9 | One-way RM ANOVA<br>Dunnett's multiple comparisons test | Treatment, F (1.492, 11.94) = 11.41 P=0.0029<br>Multiple comparisons:<br>Baseline vs. During SD P = 0.0013<br>Baseline vs. After SD P = 0.9993 |
| 1H_REM | 9 | One-way RM ANOVA<br>Dunnett's multiple comparisons test | Treatment, F (1.484, 11.87) = 5.007 P=0.0342<br>Multiple comparisons:<br>Baseline vs. During SD P = 0.1146<br>Baseline vs. After SD P = 0.9315 |
| 1I_Wake | 9 | One-way RM ANOVA<br>Dunnett's multiple comparisons test | Treatment, F (1.926, 15.40) = 10.27 P=0.0016<br>Multiple comparisons:<br>Baseline vs. During SD P = 0.0076<br>Baseline vs. After SD P = 0.9136 |
| 1I_NREM | 9 | One-way RM ANOVA<br>Dunnett's multiple comparisons test | Treatment, F (1.800, 14.40) = 7.717 P=0.0063<br>Multiple comparisons:<br>Baseline vs. During SD P = 0.0139<br>Baseline vs. After SD P = 0.9992 |
| 1I_REM | 9 | One-way RM ANOVA<br>Dunnett's multiple comparisons test | Treatment, F (1.929, 15.43) = 7.586 P=0.0054<br>Multiple comparisons:<br>Baseline vs. During SD P = 0.0068<br>Baseline vs. After SD P = 0.3908 |
| 1J | No shake: 6<br>SD 1d: 4<br>SD 5d: 6 | One-way ANOVA<br>Tukey's multiple comparisons test | Treatment, F (2, 13) = 2.012 P=0.1733<br>Multiple comparisons:<br>No shake vs. SD 1d P = 0.1990<br>No shake vs. SD 5d P = 0.2992<br>SD 1d vs. SD 5d P = 0.9003 |
| 2C | Ctrl: 7<br>SD: 6 | Two-way RM ANOVA<br>Factor 1: Interacting Object (Tria 1 E, Tria 1 S1, Tria 2 S1, Tria 2 S2)<br>Factor 2: Treatment (Ctrl, SD)<br>Tukey's multiple comparisons test | Interacting Object x Treatment, F (3, 33) = 1.824 P=0.1620<br>Interacting Object, F (3, 33) = 28.86 P<0.0001<br>Treatment, F (1, 11) = 0.009492 P=0.9241<br>Multiple comparisons P:<br>Ctrl SD<br>Trial 1 E vs. Trial 1 S1 <0.0001 0.0002<br>Trial 1 E vs. Trial 2 S1 0.998 >0.9999<br>Trial 1 E vs. Trial 2 S2 0.0086 0.9528<br>Trial 1 S1 vs. Trial 2 S1 <0.0001 0.0002<br>Trial 1 S1 vs. Trial 2 S2 0.0202 0.0007<br>Trial 2 S1 vs. Trial 2 S2 0.0054 0.9554 |
| 2D | Ctrl: 7<br>SD: 6 | Two-way RM ANOVA<br>Factor 1: Trial (Tria 1 Sociality, Tria 2 Social Novelty)<br>Factor 2: Treatment (Ctrl, SD)<br>Bonferroni's multiple comparisons test | Trial x Treatment, F (1, 11) = 3.952 P=0.0723<br>Trial, F (1, 11) = 8.775 P=0.0129<br>Treatment, F (1, 11) = 13.06 P=0.0041<br>Multiple comparisons P:<br>Ctrl vs. SD<br>Sociality >0.9999<br>Social Novelty 0.0027 |
| 2E | Ctrl: 7 | Unpaired t test with | P = 0.2800 |

| | | | |
|---|---|---|---|
| | SD: 6 | Welch's correction | |
| 2G Left | Ctrl: 6<br>SD: 5 | Two-way RM ANOVA<br>Factor 1: Trial (Tria 1, Tria 2)<br>Factor 2: Treatment (Ctrl, SD)<br>Bonferroni's multiple comparisons test | Tial x Treatment, F (1, 9) = 0.004129 P=0.9502<br>Tial, F (1, 9) = 26.67P=0.0006<br>Treatment F, (1, 9) = 0.9542 P=0.3542<br>Multiple comparisons P:<br>Trial 1 vs. Trial 2<br>Ctrl 0.0075<br>SD 0.0145 |
| 2G Right | Ctrl: 6<br>SD: 5 | Unpaired t test with Welch's correction | P = 0.7589 |
| 3A | 8 | Two-way RM ANOVA<br>Factor 1: Interacting Object (Tria 1 E, Tria 1 S1, Tria 2 S1, Tria 2 S2)<br>Factor 2: Treatment (Ctrl, SD)<br>Tukey's multiple comparisons test | Interacting Object x Treatment, F (3, 42) = 8.806 P=0.0001<br>Interacting Object, F (3, 42) = 51.86 P<0.0001<br>Treatment, F (1, 14) = 0.3361 P=0.5713<br>Multiple comparisons P:<br>Ctrl SD<br>Trial 1 E vs. Trial 1 S1 <0.0001 <0.0001<br>Trial 1 E vs. Trial 2 S1 0.9973 0.8294<br>Trial 1 E vs. Trial 2 S2 <0.0001 0.4426<br>Trial 1 S1 vs. Trial 2 S1 <0.0001 0.0004<br>Trial 1 S1 vs. Trial 2 S2 0.0002 0.0029<br>Trial 2 S1 vs. Trial 2 S2 <0.0001 0.9131 |
| 3B | 8 | Two-way RM ANOVA<br>Factor 1: Trial (Tria 1 Sociality, Tria 2 Social Novelty)<br>Factor 2: Treatment (Ctrl, SD)<br>Bonferroni's multiple comparisons test | Trial x Treatment, F (1, 14) = 1.017 P=0.3304<br>Trial, F (1, 14) = 9.823 P=0.0073<br>Treatment, F (1, 14) = 28.45 P=0.0001<br>Multiple comparisons P:<br>Ctrl vs. SD<br>Sociality 0.0073<br>Social Novelty 0.0002 |
| 3C | 6 | Two-way RM ANOVA<br>Factor 1: Interacting Object (Tria 1 E, Tria 1 S1, Tria 2 S1, Tria 2 S2)<br>Factor 2: Treatment (Ctrl, SD)<br>Tukey's multiple comparisons test | Interacting Object x Treatment, F (3, 30) = 3.359 P=0.0317<br>Interacting Object, F (3, 30) = 57.33 P<0.0001<br>Treatment, F (1, 10) = 3.184 P=0.1047<br>Multiple comparisons P:<br>Ctrl SD<br>Trial 1 E vs. Trial 1 S1 <0.0001 <0.0001<br>Trial 1 E vs. Trial 2 S1 >0.9999 0.9046<br>Trial 1 E vs. Trial 2 S2 <0.0001 <0.0001<br>Trial 1 S1 vs. Trial 2 S1 <0.0001 <0.0001<br>Trial 1 S1 vs. Trial 2 S2 0.9972 0.0209<br>Trial 2 S1 vs. Trial 2 S2 <0.0001 <0.0001 |
| 3D | 6 | Two-way RM ANOVA<br>Factor 1: Trial (Tria 1 Sociality, Tria 2 Social Novelty)<br>Factor 2: Treatment (Ctrl, SD)<br>Bonferroni's multiple comparisons test | Tial x Treatment, F (1, 10) = 2.302 P=0.1602<br>Tial, F (1, 10) = 1.955 P=0.1923<br>Treatment, F (1, 10) = 0.6255 P=0.4474<br>Multiple comparisons P:<br>Ctrl vs. SD<br>Sociality 0.2208<br>Social Novelty >0.9999 |
| 4A | WT: 9 | Two-way RM ANOVA | Interacting Object x Genotype, F (3, 45) = 1.971 P=0.1319 |

| | | | |
|---|---|---|---|
| | Homo: 8 | Factor 1: Interacting Object (Tria 1 E, Tria 1 S1, Tria 2 S1, Tria 2 S2)<br>Factor 2: Genotype (WT, Homo)<br>Tukey's multiple comparisons test | Interacting Object, F (2.394, 35.91) = 38.98 P<0.0001<br>Genotype, F (1, 15) = 0.6145 P=0.4453<br>Multiple comparisons P:<br>WT / Homo<br>Trial 1 E vs. Trial 1 S1: <0.0001 / 0.0038<br>Trial 1 E vs. Trial 2 S1: 0.9997 / 0.2193<br>Trial 1 E vs. Trial 2 S2: 0.037 / 0.4328<br>Trial 1 S1 vs. Trial 2 S1: <0.0001 / 0.0535<br>Trial 1 S1 vs. Trial 2 S2: 0.006 / 0.083<br>Trial 2 S1 vs. Trial 2 S2: 0.0116 / 0.9801 |
| 4B | WT: 5<br>Homo: 6 | Unpaired t test with Welch's correction | Wake, P = 0.0231<br>NREM, P = 0.0895<br>REM, P = 0.0308 |
| 4C | WT: 5<br>Homo: 6 | Unpaired t test with Welch's correction | Wake, P = 0.7179<br>NREM, P = 0.6378<br>REM, P = 0.6168 |
| 4E | 12 (WT 4, Het 3, Homo 5) | Simple linear regression | Sociality / Social Novelty<br>Goodness of Fit<br>R squared: 0.4714 / 0.007258<br>Is slope significantly non-zero?<br>F: 8.916 / 0.07311<br>DFn, DFd: 1, 10 / 1, 10<br>P value: 0.0137 / 0.7924 |
| 4F | 12 (WT 4, Het 3, Homo 5) | Simple linear regression | Sociality / Social Novelty<br>Goodness of Fit<br>R squared: 0.01864 / 0.4250<br>Is slope significantly non-zero?<br>F: 0.1899 / 7.390<br>DFn, DFd: 1, 10 / 1, 10<br>P value: 0.6722 / 0.0216 |
